# A synaptic mechanism for encoding the learned value of action-derived safety

**DOI:** 10.1101/2025.08.04.668532

**Authors:** Emma E. Macdonald, Jun Ma, Di Liu, Kai Yu, Michael E. Authement, Yan Leng, Hannah C. Goldbach, Veronica A. Alvarez, Bruno B. Averbeck, Mario A. Penzo

## Abstract

Motivated behavior is often framed in terms of biologically grounded outcomes, such as food or threat. Yet many motivated actions, like the pursuit of safety or agency, depend on outcomes that lack explicit sensory value and must instead be inferred from experience. Here, we identify a thalamostriatal circuit mechanism by which such internally constructed outcomes acquire motivational value. In mice performing an active avoidance task, neurons in the paraventricular thalamus (PVT) projecting to the nucleus accumbens (NAc) develop a safety-encoding signal that emerges following successful avoidance. This signal is experience-dependent and value-sensitive, diminishing upon devaluation of the instrumental contingency. Selective silencing of the PVT→NAc projection at safety onset disrupts avoidance persistence without impairing action-outcome learning, as confirmed by computational modeling of value updating based on prediction error. Mechanistically, PVT input recruits cholinergic interneurons (CINs) to modulate dopamine release and this influence depends on synaptic potentiation mediated by GluA2-lacking AMPA receptor insertion at PVT–CIN synapses. Disrupting this plasticity selectively impairs the avoidance response by blunting the motivational value of safety without affecting acquisition. These findings reveal how thalamic circuits assign value to abstract, internal outcomes, providing a framework for understanding how goals like safety are inferred, stabilized, and rendered behaviorally effective.

## MAIN

Many adaptive behaviors unfold in the absence of primary reinforcers. While the neuroscience of motivation has largely centered on the pursuit of innate rewards and the avoidance of sensory threats, a substantial portion of behavioral control is directed toward internal goals, such as safety, predictability, or cognitive resolution, that lack intrinsic sensory value and must be constructed through experience (1). These goals are not hardwired but inferred, and their motivational value must be learned, represented, and stabilized internally. Despite their central role in both human and animal behavior, the neural mechanisms by which such non-primary outcomes acquire value and sustain future actions remain poorly understood.

A prominent example of such a constructed goal is safety: an internal state inferred from the successful avoidance of harm. Unlike passive safety cues, which signal threat omission without requiring behavior, instrumentally generated safety, as observed in active avoidance, generates a qualitatively distinct motivational signal (2–4). This form of safety promotes behavioral persistence, reinforces future avoidance, and evokes dopaminergic responses analogous to those observed during reward acquisition (5–12). While the contribution of corticostriatal circuits to value encoding is well established, the potential role of thalamic circuits in shaping the learned value of internal outcomes has received comparatively little attention. Whether thalamic inputs can support the internal construction and stabilization of motivational value, and how such representations interact with dopaminergic systems, remains largely unexplored.

To investigate how the brain assigns value to internal, non-sensory outcomes, we leveraged an active avoidance paradigm in which animals learn to escape an impending threat. This task provides a behavioral window into the construction of safety as an inferred goal state. We focused on the paraventricular thalamus (PVT), a dorsal midline structure positioned to integrate interoceptive and motivational signals and known to project densely to the nucleus accumbens (NAc), a key node in reward and motivational circuits (13–15). Prior work has suggested that the PVT may relay internal drive states and contribute to motivational salience (16–18) and valence assignment (19), yet whether it participates directly in encoding outcome value signals that support goal-oriented behavior has remained unclear. We combined pathway-specific optogenetic perturbations, fiber photometry, synaptic physiology, and computational modeling to test whether PVT→NAc projections are necessary and sufficient for encoding the motivational value of successful avoidance. Our results uncover an experience-dependent plastic interaction between PVT inputs and striatal cholinergic interneurons that modulates dopaminergic tone and is necessary for maintaining the learned value of safety, but not the acquisition of instrumental actions. These findings point to an unexpected role for the thalamus in stabilizing internal outcome representations that sustain adaptive behavior.

### PVT→NAc projections encode the learned value of action-derived safety

To determine how internally inferred outcomes such as safety acquire motivational value, we trained mice in a two-way active avoidance (2AA) task, in which they learned to shuttle across a hurdle during a warning signal (WS) to terminate the WS and avoid a predicted footshock (See Methods). Following a successful avoidance shuttle, action-generated stimuli that are negatively correlated with shock, such as WS offset, are thought to reinforce avoidance behavior by serving as inferred safety signal (20). We recorded calcium dynamics from GCaMP8s-expressing terminals of PVT neurons in the NAc during training (days 1, 3, and 5) to track the emergence of PVT→NAc activity (Fig. 1A–B; S1). Population heatmaps aligned to WS onset and sorted by escape latency revealed a continuum of activity changes that spanned the cue period and increased with training (Fig. 1C,D). However, inspection of individual trials (e.g., Fig. 1E) suggested more discrete increases in PVT→NAc activity time-locked to specific behavioral events, namely shuttle initiation and the onset of inferred safety, as seen in previous studies (21). Aligning activity to these discrete events confirmed distinct peaks at both time points (Fig. 1F,G). We focused on the safety-related calcium signal, which suggests that PVT→NAc projections become engaged during the inferred experience of having successfully avoided harm.

**Figure 1:**
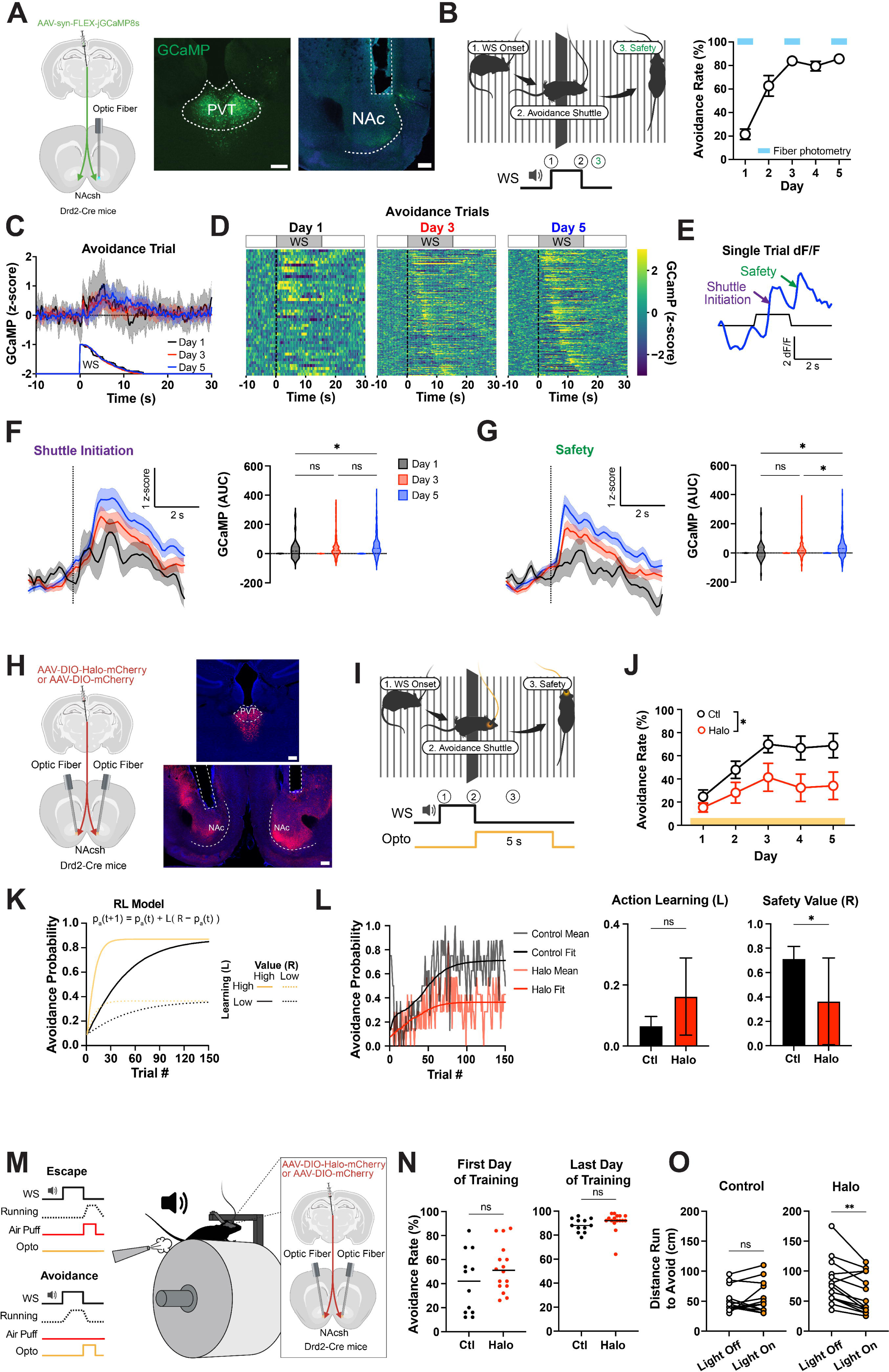
pPVT→NAc projections encode the value of inferred safety and sustain motivation during avoidance. **(A)** Drd2-Cre mice were injected with GCaMP8s in pPVT to record terminal activity in NAc (left). [Art created in Biorender]. Representative images of injection and recording sites (right). Scale bars, 200μm. **(B)** Schematic of behavior apparatus and example avoidance trial depicting behavior sequence at WS onset (left). Performance in the 2AA task (n=7 mice) (right). **(C)** Heatmaps of calcium signals during avoidance trials, ordered by avoidance latency (low to high). **(D)** Representative calcium dF/F signal during avoidance trial. **(E)** GCaMP dF/F z-score normalized in all avoidance trials by day. **(F)** GCaMP dF/F z-score normalized at shuttle initiation (dotted line) in avoidance trials (left). Quantification of post event activity (right). AUC, pairwise comparison across days, mixed effects model: see statistics table. **(G)** GCaMP dF/F z-score normalized at safety (dotted line) in avoidance trials (left). Quantification of post event activity (right). AUC, pairwise comparison across days, mixed effects model: see statistics table. **(H)** Drd2-Cre mice were injected with halorhodopsin (Halo) or fluorescent control in pPVT to inhibit terminal activity in NAc (left). Representative images of injection and stimulation sites (right). Scale bars, 200μm. **(I)** Schematic of closed-loop optogenetic manipulation in the 2AA task (see Methods). **(J)** Performance in the 2AA task with inhibition of pPVT-NAc at safety (Control n=8, Halo n=7 mice). 2-way Anova group effect: p=0.0345. **(K)** Simulated avoidance probability displaying the effect of changes to learning and value parameters in learning model (pa = probability of avoidance, R = value of safety, L = learning rate). **(L)** Avoidance probability across trial with best fit model probability by group (left). Best fit learning parameter by group with bootstrap estimated SEM. Bootstrap permutation test for group difference: p=0.1032 (middle). Best fit value parameter by group with bootstrap estimated SEM. Bootstrap permutation test for group difference: p=0.0448 (right). **(M)** Schematic of head-fixed avoidance task and injection targets (see Methods). **(N)** Average behavior performance avoidance training with no optogenetic stimulation on the first (left, Unpaired t test: p=0.1771) and last (right, Unpaired t test: p=0.2493) days of training. Control n=13, Halo n=16 mice. **(O)** Sessions breakpoints for progressive ratio test (5 cm increments) for Halo (left, Paired t test: p=0.0035) and mCherry (right, Paired t test: p=0.4947) groups. Data are shown as mean ± s.e.m. *p < 0.05, **p < 0.01, ns: p > 0.05.

To test the functional relevance of this signal, we used closed-loop optogenetic silencing of PVT→NAc terminals immediately after successful avoidance. Across days, inhibition of this pathway impaired the acquisition of avoidance behavior relative to controls (Fig. 1H–J, S1C). To probe the underlying computational mechanism and dissociate learning of action-outcome associations (action learning) and safety value assignment to outcome (value learning) we applied a reinforcement learning model wherein action learning and value learning were estimated separately (Fig. 1K, Methods). This analysis revealed that behavioral deficits following PVT→NAc inhibition were best explained by a selective reduction in the value of the safety outcome, rather than a change in action learning rate (Fig. 1L). Conversely, optogenetic excitation of PVT→NAc terminals during the safety period enhanced modeled outcome value and suppressed action learning rate (Fig. S2), which suggests that excessive activation has less specific effects than inactivation. Fitting a bilinear model to the excitation data confirmed a significant upward shift in intercept without a change in slope, reinforcing the conclusion that this projection encodes outcome value (Fig. S2F-G).

We next asked whether this safety signal was sensitive to devaluation of instrumental contingency, which is an important criterion for determining whether a neural signal encodes value rather than reflecting a fixed stimulus–response association. Trained animals underwent an extinction-with-response-prevention session in which a transparent barrier blocked shuttling, and the WS no longer predicted shock. When tested in subsequent extinction sessions (with the barrier removed and no shock delivered), mice showed an approximately 50% reduction in avoidance behavior. This behavioral change was accompanied by a significant decrease in the safety-evoked PVT signal (Fig. S3A–D). When the shock contingency was reinstated, both avoidance behavior and PVT→NAc safety-related activity rebounded, often surpassing pre-devaluation levels (Fig. S3E). This recovery further supports the interpretation that extinction-related reductions were due to value updating and not attributable to photobleaching or signal degradation.

Finally, we tested whether this thalamic derived value signal supports effortful avoidance behavior. Parametrizing effort in shuttle-based avoidance is challenging, since animals either cross or fail to cross without a clear gradient of work required. To overcome this limitation, we used a head-fixed version of the task in which mice learned to run a fixed distance to avoid an air puff (Fig. 1M). In this setting, within-subject silencing of PVT→NAc terminals reduced the break point in a progressive ratio test (See Methods), indicating a diminished willingness to work for safety (Fig. 1N-O). Together, these results demonstrate that PVT→NAc projections encode a safety-related value signal that emerges with learning of the task and sustains motivated behavior despite increased effort demands.

### PVT→NAc projections are required for safety-evoked dopamine release

Previous work has shown that dopamine (DA) levels in the NAc increase in response to safety cues during avoidance behavior, suggesting that threat omission can acquire reward-like salience (7–9,22,23). We asked whether a similar dopaminergic safety signal is present in our 2AA task and whether it depends on PVT input. To test this, we expressed the genetically encoded DA sensor dLight1.2 (24) in the NAc and recorded DA dynamics during training. We observed an increase in DA release at the onset of the safety period (WS offset) across training days (Fig. 2A– F; S4), consistent with the acquisition of motivational value. To determine whether this signal requires PVT input, we used unilateral closed-loop optogenetic silencing of PVT→NAc terminals at safety onset during dLight recordings (Fig. 2G). This manipulation did not affect behavior, confirming that the dopaminergic measurements were not confounded by performance (Fig. 2I). However, PVT silencing significantly attenuated safety-evoked DA release (Fig. 2J,K), indicating that the emergence of a dopaminergic safety signal depends on input from the PVT.

**Figure 2:**
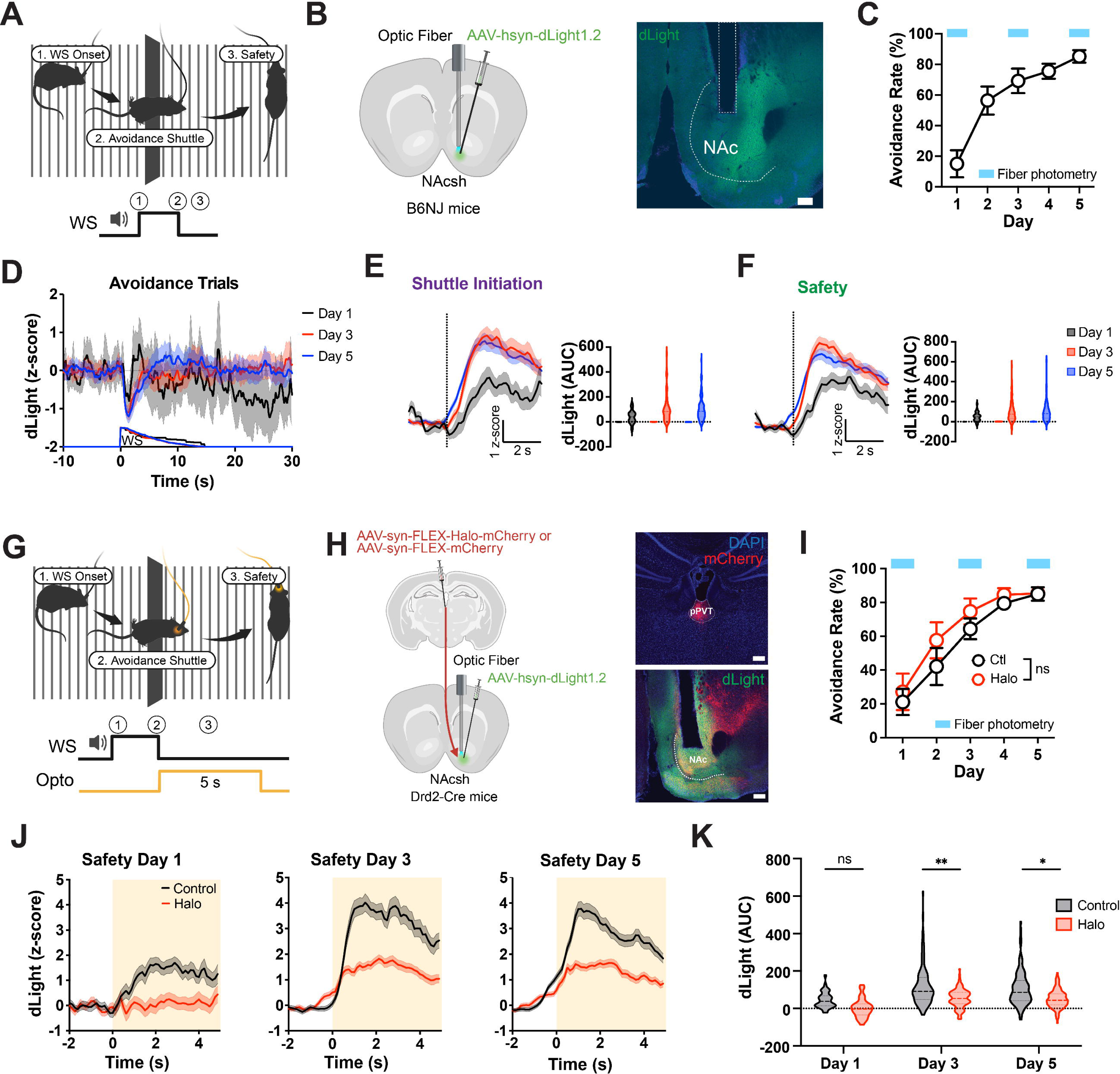
pPVT input drives NAc dopamine signals associated with safety. **(A)** Schematic of behavior apparatus and example avoidance trial depicting behavior sequence at WS onset. **(B)** Wild type mice were injected with dLight1.2 to record dopamine in NAc (left). Representative image of recording site (right). Scale bar, 200μm. **(C)** Performance in the 2AA task (n=6 mice). **(D)** dLight signal in all avoidance trials by day. **(E)** dLight dF/F z-score normalized at shuttle initiation (dotted line) in avoidance trials (left). Quantification of post event activity (right). AUC, pairwise comparison across days, mixed effects model: see statistics table. **(F)** dF/F z-score normalized at safety (dotted line) in avoidance trials (left). Quantification of post event activity (right). AUC, pairwise comparison across days, mixed effects model: see statistics table. **(G)** Schematic of closed-loop optogenetic manipulation in the 2AA task (see Methods). **(H)** Drd2-Cre mice were injected with halorhodopsin (Halo) or fluorescent control in pPVT and dLight1.2 in NAc to test the effects of pPVT-NAc inhibition of safety dopamine release (left). Representative images of injection and recording sites (right). Scale bars, 200μm. **(I)** Performance in the 2AA task. 2-way Anova: p=0.517 (Control n=6, Halo n=5 mice). **(J)** dLight dF/F z-score normalized at safety with optogenetic inhibition by group day 1 (left), day 3 (middle), and day 5 (right) of 2AA training. **(K)** Quantification of post event activity. AUC, pairwise comparisons between groups, linear mixed-effects model for repeated measures: see statistics table. Data are shown as mean ± s.e.m. *p < 0.05, **p < 0.01, ns: p > 0.05.

### Cholinergic interneurons mediate PVT-evoked dopamine and are increasingly recruited during safety value learning

While PVT projections are necessary for safety-related DA release, prior anatomical studies suggest that midline thalamic inputs, including those from PVT, do not directly innervate dopaminergic terminals in the NAc (25). Instead, they synapse heavily onto striatal cholinergic interneurons (CINs), which have been shown to regulate DA release through nicotinic mechanisms (26–33). Importantly, recent work from Baimel et al. (2022) demonstrated that PVT is a major source of input to NAc CINs, providing a potential anatomical substrate for PVT-dependent modulation of DA. To directly test this, we first used fast-scan cyclic voltammetry (FSCV) in acute slices to measure DA release evoked by PVT terminal stimulation. Application of the nicotinic antagonist DHβE abolished the DA response (Fig. 3A-C), indicating that PVT-evoked DA release requires cholinergic transmission. We next asked whether CINs themselves are recruited by safety value learning. Using the ACh sensor gACh3.8 (34), we recorded cholinergic activity in the NAc across 2AA training. The onset of the safety period was associated with a phasic ACh response that increased with training (Fig. 3D-J; S4), mirroring the dynamics observed in PVT and DA signals. Finally, we assessed whether CIN activity is required for PVT-evoked DA release *in vivo*. To do this, we expressed dLight1.2 in the NAc of ChAT-Cre mice, optogenetically stimulated PVT terminals, and simultaneously silenced CINs using the *<u>D</u>*esigner *<u>R</u>*eceptors *<u>E</u>*xclusively *<u>A</u>*ctivated by *<u>D</u>*esigners *<u>D</u>*rugs (DREADDs) hM4Di. Interestingly, PVT-evoked DA was potentiated by 2AA training, suggesting that avoidance learning induces plasticity within this circuit. However, chemogenetic inhibition of CINs completely abolished PVT-evoked DA release following training (Fig. 3K-O), providing convergent evidence for a PVT→CIN→DA pathway. Together, these findings support a model in which safety learning enhances the ability of PVT inputs to engage local downstream dopaminergic signaling through cholinergic interneurons.

**Figure 3:**
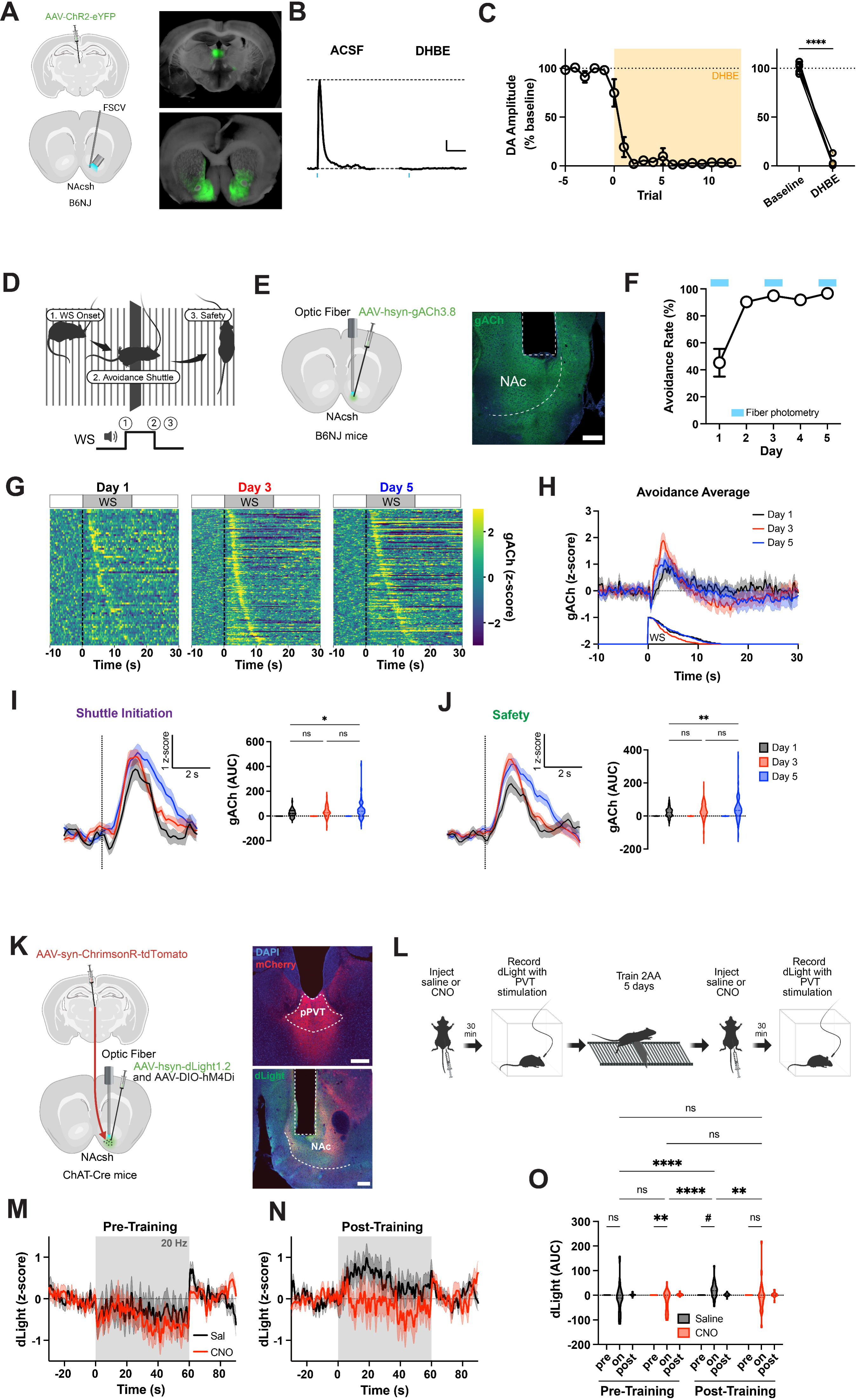
NAc acetylcholine is necessary for pPVT-driven dopamine signaling after safety learning. **(A)** Wildtype mice were injected with channelrhodopsin in pPVT and fast-scan cyclic voltammetry was used on NAc slices to measure optogenetically evoked dopamine (right). Representative images of injection and recording sites with ChR-eYP expression (right). **(B)** Representative PVT evoked [DA] in ACSF and DHBE. Scale, 100nM, 2s. **(C)** oDA amplitude normalized to baseline in ACSF and DHBE (left). Average oDA amplitude of three trials by slice (right). Paired t test: p<0.0001 (n=5 slices from 5 mice). **(D)** Schematic of behavior apparatus and example avoidance trial depicting behavior sequence at WS onset. **(E)** Wildtype mice were injected with GRAB ACh3.8 (gACh) in NAc to record ACh activity in 2AA (left). Representative image of recording site (left). Scale bar, 200um. **(F)** Performance in the 2AA task (n=5 mice). **(G)** Heatmaps of gACh signals during avoidance trials, ordered by avoidance latency (low to high). **(H)** gACh signal in all avoidance trials by day. **(I)** gACh dF/F z-score normalized at shuttle initiation (dotted line) in avoidance trials (left). Quantification of post event activity (right). AUC, pairwise comparison across days, mixed effects model: see statistics table. **(J)** gACh dF/F z-score normalized at safety (dotted line) in avoidance trials (left). Quantification of post event activity (right). AUC, pairwise comparison across days, mixed effects model: see statistics table. **(K)** ChAT-Cre mice were injected with ChrimsonR in pPVT and dLight1.2 and DIO-hM4Di in NAc (left). Representative images of injection and recording sites (right). Scale bar, 200um. **(L)** Schematic of recording and training schedule (see Methods). **(M-N)** dLight during pPVT terminal stimulation by treatment pre-training (**M**) and post-training (**N**). **(O)** Quantification of dLight activity. AUC, pairwise comparisons, mixed effects model: see statistics table (n=4 mice). Data are shown as mean ± s.e.m. *p < 0.05, **p < 0.01, ****p < 0.0001, ns: p > 0.05, #: p<0.08.

### Safety value learning induces synaptic plasticity at PVT→CIN synapses

The potentiation of PVT-evoked, CIN-dependent DA release in NAc suggests experience-dependent plasticity within the PVT→CIN microcircuit. We next asked whether this functional enhancement arises from synaptic modifications at PVT→CIN conenctions following avoidance training. To test this, we injected ChAT-Cre mice with a Cre-dependent mCherry reporter in the NAc to label CINs and expressed ChR2-eYFP in PVT neurons to enable selective optical activation of PVT→NAc terminals (Fig. 4A). Mice were then trained in the 2AA task or assigned to a yoked control group that received identical shock and cue exposure, but without control over shock omission (Fig. 4B). As expected, only trained animals developed robust avoidance behavior by day 5, while yoked animals did not develop shuttle responses to the WS (Fig. 4C). Two hours after the final training session (Day 5), acute coronal brain slices containing the NAc were prepared, and we recorded optogenetically evoked excitatory postsynaptic currents (oEPSCs) in fluorescently identified CINs. Compared to yoked controls, trained animals exhibited significantly larger AMPA-mediated oEPSCs (Fig. 4D,E) along with an elevated AMPA/NMDA ratio (Fig. 4I). NMDA current amplitudes were not significantly different between groups (Fig. 4H), suggesting a postsynaptic locus of plasticity. However, a significant reduction in the paired-pulse ratio (PPR) was also observed in trained mice (Fig. 4J), indicating a possible contribution of presynaptic release probability. We next asked whether postsynaptic receptor composition had changed following training. The kinetics of AMPA currents in trained mice showed significantly faster rise and decay times relative to yoked controls (Fig. 4F,G), consistent with insertion of GluA2-lacking, calcium-permeable AMPA receptors (35–37). Supporting this interpretation, rectification indices were significantly higher in CINs from trained animals, and AMPA currents were more strongly attenuated by NASPM, a selective blocker of GluA2-lacking AMPARs (Fig. 4K,L). Together, these data demonstrate that active avoidance training induces synaptic potentiation at PVT→CIN synapses in the NAc, characterized by enhanced AMPA receptor transmission and recruitment of GluA2-lacking AMPARs. This plasticity likely underlies the increased ability of PVT input to drive DA observed after training and provides a plausible mechanistic substrate for the modulation of safety-related dopamine signaling.

**Figure 4:**
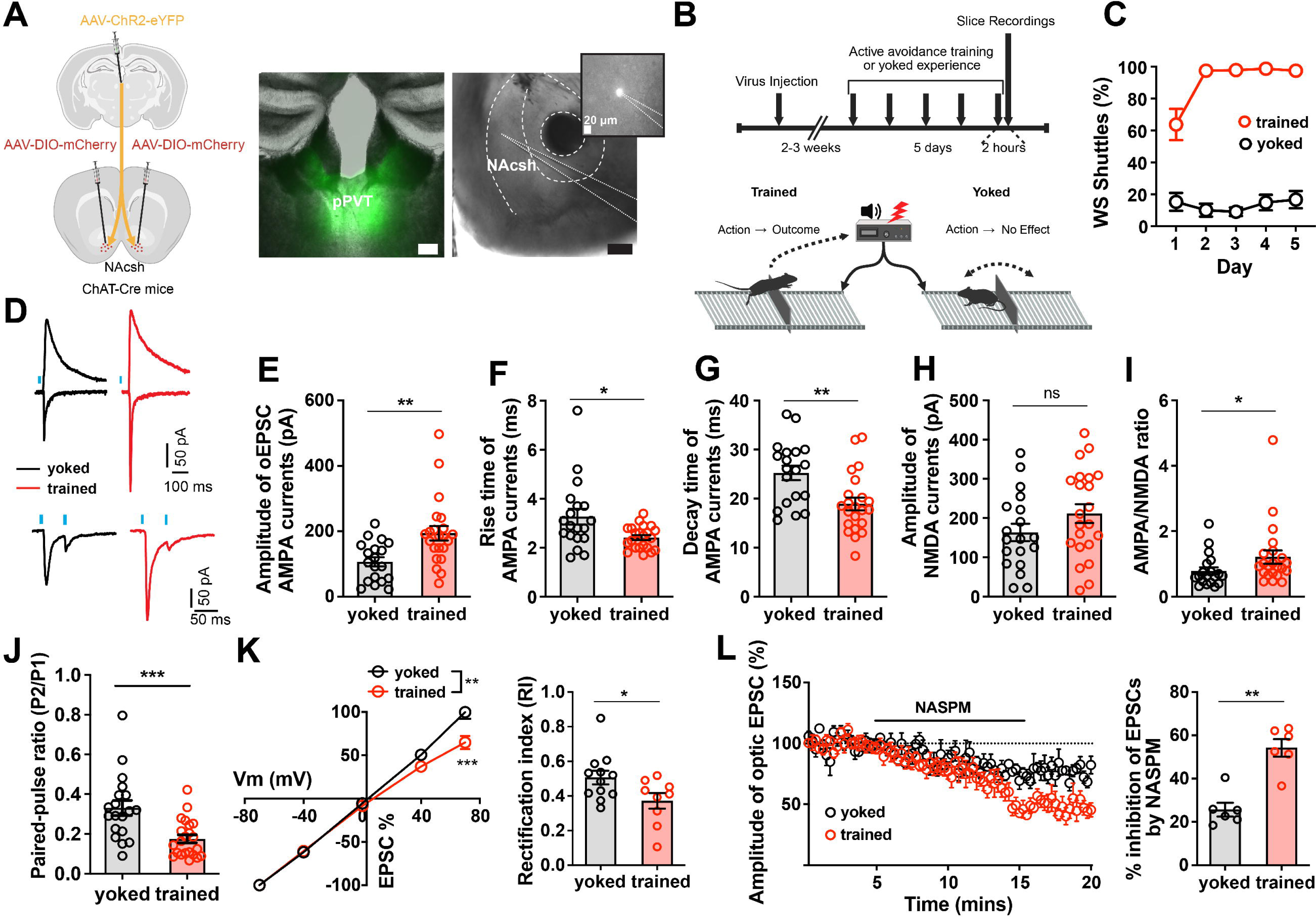
Active avoidance training induces plasticity at pPVT→CIN synapses. **(A)** ChAT-Cre mice were injected with channelrhodopsin in pPVT and DIO-mCherry in NAc to record post-synaptic currents in CINs with pPVT stimulation (left). Representative images of injection and recording sites (right). Scale bars, 200um. **(B)** Schematic of training and recording schedule (see Methods). **(C)** Performance in the 2AA task (trained n= 8, yoked n=7 mice). **(D)** Representative traces of optogenetic evoked voltage clamp responses. **(E-J)** Measurements of current properties. Unpaired t tests: oEPSC amplitude p=0.0024; Rise time p=0.0107; Decay time p=0.0028; NMDA amplitude p=0.1562; Paired-pulse ratio p=0.0005. Mann Whitney test: AMPA/NMDA ratio p=0.0168 (trained n=22 neurons from 4 mice, yoked n=19 neurons from 3 mice). **(K)** Rectification curve (left). 2way Anova: p=0.0085. Rectification index (right). Unpaired t test: p=0.0360. **(L)** Amplitude of evoked response with NASPM application (left). Quantification of NASPM inhibition (right). Unpaired t test: p=0.0002 (trained n=9 neurons from 4 mice, yoked n=12 neurons from 4 mice). Data are shown as mean ± s.e.m. *p < 0.05, **p < 0.01, ***p < 0.001, ns: p > 0.05.

### GluA1-dependent plasticity at PVT→CIN synapses is necessary for encoding the value of safety

The previous results demonstrated that avoidance training potentiates PVT→CIN synapses through the recruitment of GluA2-lacking AMPA receptors, a form of plasticity classically associated with GluA1 homomeric receptor insertion (38,39). To test whether GluA1 is necessary for this synaptic remodeling, we used a CRISPR-Cas9 strategy to selectively delete Gria1, the gene encoding GluA1, from cholinergic interneurons. We injected ChAT-Cre mice unilaterally with a Cre-dependent CRISPR construct (AAV9-EF1α-DIO-Cas9-sgGria1), which co-expresses Cas9 and a guide RNA targeting *Gria1* (40) and injected the contralateral hemisphere with a control virus lacking the sgRNA (Cas9-empty) (Fig. 5A). Mice were then trained in the 2AA task, and brain slices were prepared two hours after the final training session for whole-cell recordings from CINs in each hemisphere. In the control (Cas9-empty) hemisphere, CINs displayed properties consistent with synaptic potentiation as previously observed: elevated AMPA oEPSCs, increased AMPA/NMDA ratios, greater rectification, and enhanced sensitivity to NASPM (Fig. 5D-K). In contrast, CINs in the sgGria1-injected hemisphere exhibited significantly reduced AMPA-mediated currents, lower AMPA/NMDA ratios, and decreased rectification indices (Figure 5D-K). NASPM sensitivity was also attenuated, consistent with a failure to incorporate GluA2-lacking AMPA receptors in the absence of GluA1 (Fig. 5L,M). These findings confirm that GluA1 expression in CINs is required for the experience-dependent synaptic plasticity induced by safety value learning.

**Figure 5:**
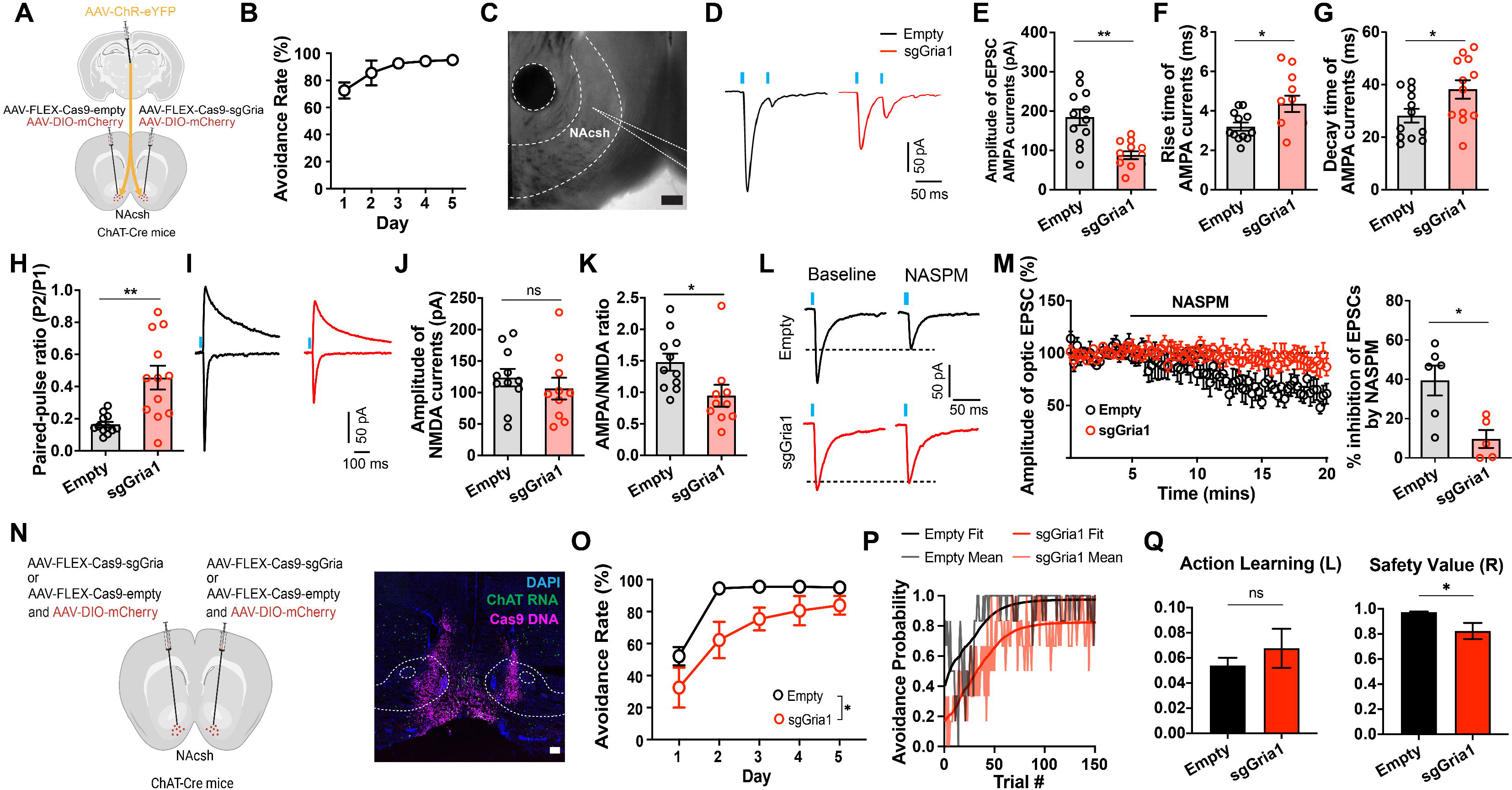
NAc CIN Gria1 knockout disrupts avoidance-related plasticity and safety valuation. **(A)** ChAT-Cre mice were injected with channelrhodopsin in pPVT, and FLEX-Cas9-sgGria1 and empty control contralaterally in NAc to record post-synaptic currents in CINs with pPVT stimulation (left). **(B)** Performance in the 2AA task (n=5 mice). **(C)** Representative image of recording location. Scale bar, 200um. **(D)** Representative oEPSCs from control and knockout hemispheres. **(E-H)** Measurements of current properties. Unpaired t tests: oEPSC amplitude p=0.0003; Rise time p=0.0206; Decay time p=0.0321; Paired-pulse ratio p=0.0009. **(I)** Representative traces of AMPA and NMDA responses to pPVT optogenetic stimulation. **(J-K)** Current amplitude measurements. Unpaired t tests: NMDA amplitude p=0.4374; AMPA/NMDA ratio p=0.0256 (sgGria1 n=12 neurons, empty n=12 neurons from 4 mice). **(L)** Representative traces of voltage clamp responses to optogenetic stimulation before and after NASPM application. **(M)** Amplitude of evoked response with NASPM application (left). Quantification of NASPM inhibition (right). Unpaired t test: p=0.0113 (sgGria1 n=5 neurons, empty n=6 neurons from 5 mice). **(N)** ChAT-Cre mice were injected with FLEX-Cas9-sgGria or control bilaterally in NAc (left). Representative image of virus spread through fluorescent in situ hybridization Cas9 probe binding to viral DNA (magenta) and ChAT mRNA expression (green) (right). **(O)** Performance in the 2AA task by group. Mixed-effects group effect: p=0.0478 (sgGria1 n=6, empty n=6 mice). **(P)** Avoidance probability across trial with best fit model probability by group (left). Best fit parameters by group with bootstrap estimated error. Bootstrap permutation tests for group difference: Learning p=0.2136; Value p=0.0372. Data are shown as mean ± s.e.m. *p < 0.05, **p < 0.01, ns: p > 0.05.

While post-shuttle PVT→NAc signaling reflects the inferred value of avoidance-derived safety outcomes, in theory, synaptic plasticity at PVT→CIN synapses could either support action or value learning. To probe these two possibilities, we bilaterally injected ChAT-Cre mice with the *Gria1*-targeting CRISPR construct and trained them in the 2AA task. Compared to control animals, CRISPR-mediated deletion of GluA1 in CINs significantly impaired the acquisition of avoidance behavior (Fig. 5N,O; S6). To assess whether this deficit reflected a disruption in outcome valuation versus action learning, we applied RL modeling to trial-by-trial behavior. Strikingly, GluA1 deletion selectively impaired the value parameter while leaving the learning rate of avoidance intact (Fig. 5P,Q), mirroring the effects of PVT→NAc silencing and directly implicating CIN plasticity as a critical node for encoding internal safety value. Together, these findings identify GluA1-dependent synaptic plasticity at PVT→CIN synapses as a key mechanism by which safety value learning is integrated into striatal circuits to support motivational drive. This local plasticity enables thalamic input to more effectively engage cholinergic modulation and downstream dopamine signaling, stabilizing the inferred value of threat omission and sustaining safety-seeking behavior.

## Discussion

Many goals that guide behavior lack inherent sensory value and must be internally inferred (1). Our findings reveal a thalamostriatal mechanism by which such abstract, action-contingent outcomes acquire motivational weight. This extends classical models of valuation, which emphasize external cues and innate reinforcers (13,41) by demonstrating that the PVT can endow internally generated outcomes with behavioral utility.

Our findings support a conceptualization of the PVT as part of a central system for prioritizing internally constructed goals and broadcasting value-based signals to guide behavior. Depending on the target, this broadcast can serve distinct computational functions. For instance, PVT projections to the amygdala have been shown to assign motivational significance to sensory events, shaping cue–outcome associations (19). In contrast, the PVT→NAc pathway described here appears to support the stabilization of internal goal representations by assigning value to internally inferred outcomes, thereby promoting the actions required to obtain them. This dual capacity—assigning value to external cues and elevating the value of internally generated outcomes—positions the PVT as a key node in the translation of internal demands into motivational control. As such, our results support and extend the emerging view that the thalamus is not a passive relay, but an active selector of behaviorally relevant information serving not only as an outward-facing searchlight that prioritizes external stimuli for perceptual awareness (42–45), but also as an inward-facing one that elevates internal goals for motivational control (17).

PVT’s role in prioritizing internal goals is further shaped by physiological and emotional states. Prior studies have shown that PVT engagement enhances the attractiveness of food odors during hunger (46) and suppresses reward-seeking under threat (47–49), highlighting its function as a dynamic gate that adjusts motivational salience based on internal state (50–52). Our findings build on this framework by identifying a synaptic mechanism through which PVT signals relays the motivational value of an inferred, action-contingent outcome. This provides mechanistic insight into how internal states influence not only cue salience, but also the motivational strength of internal goals.

At the circuit level, we uncover a disynaptic architecture in which thalamic input engages dopaminergic signaling through modulation of CINs in the striatum. CINs are well known to regulate DA release (29,30,53,54) and contribute to motivated behavior (55,56). Recent studies further show that CINs are required for effortful reward pursuit (57) and that glutamatergic plasticity onto CINs shapes reinforcement-driven decision-making (58). Our findings add a new dimension to this literature by showing that synaptic strengthening at PVT→CIN synapses increases CIN recruitment and amplifies dopaminergic output in response to an internally generated goal. GluA1-dependent incorporation of calcium-permeable AMPARs at these synapses may serve as a local gain mechanism for motivational drive.

These results do not suggest that CINs are exclusively dedicated to inferred outcomes, but rather that their engagement depends on the identity and plasticity of upstream input. In this case, PVT input undergoes experience-dependent strengthening, rendering the circuit more effective at coupling internal signals to dopaminergic modulation. CINs thus emerge as flexible integrators of excitatory input whose contribution to behavior is shaped by the structure of learning. More broadly, our findings support a model in which valuation emerges from interactions across distributed hierarchies: thalamic input stabilizes goal representations, CINs provide amplification, and dopamine modulates effort and learning accordingly (59).

This architecture may be critical for sustaining goal-directed behavior in the absence of immediate sensory incentives. Dysregulation of thalamostriatal plasticity could distort the motivational value of internal goals, either by inflating their perceived importance, as may occur in pathological avoidance, or by attenuating their salience, as observed in the amotivation and avolition associated with schizophrenia (60–63). Postmortem studies have revealed reduced expression of cholinergic interneuron markers in the striatum of individuals with schizophrenia, suggesting a link between CIN dysfunction and impaired motivational control (64). Recent work in a genetic mouse model of schizophrenia has further established a mechanistic link between striatal cholinergic dysfunction, impaired thalamostriatal transmission, and the emergence of an avolition-like phenotype (61). By linking synaptic plasticity to motivational drive, our findings offer a mechanistic foundation for understanding how internal outcomes are prioritized and how such prioritization may become distorted in psychiatric disease.

Looking ahead, an important next step will be to determine whether similar thalamostriatal mechanisms support other forms of internally generated value, such as reducing uncertainty, resolving cognitive conflict, experiencing relief, or satisfying curiosity. It will also be important to investigate how such value signals are shaped by interoceptive states like hunger, arousal, or stress, and how these interactions influence behavioral prioritization. More broadly, these findings raise new questions about how the PVT coordinates with cortical and limbic systems to arbitrate between internal goals and external demands. By identifying a circuit mechanism through which abstract action-contingent outcomes gain motivational significance, this work advances our understanding adaptive of how the brain constructs and updates behavioral priorities in dynamic environments.

## Materials and Methods

### Subjects

All procedures were performed in accordance with the Guide for the Care and Use of Laboratory Animals and were approved by the National Institute of Mental Health (NIMH) Animal Care and Use Committee. Mice used in this study were group housed under a 12-h light/dark cycle (6:00– 18:00 light), at temperatures of 70–74 °F and 40–65% humidity, with food and water available *ad libitum*. After surgery, mice were singly housed. C57BL/6NJ strain mice (The Jackson Laboratory #005304) were crossed with Drd2-Cre (GENSAT founder line ER44), ChAT-Cre (The Jackson Laboratory # 018957), or DAT-Cre (#006660). Both male and female mice 8–20 weeks of age were used for all experiments. Animals were randomly allocated to the different experimental conditions reported in this study.

### Viral vectors

AAV9-hSyn-Flex-GCaMP8s-WPRE (#162377), pAAV.Syn.Flex.GCaMP6s.WPRE.SV40 (#100845), pAAV-hSyn-dLight1.2 (#111068), pAAV-hsyn-GRAB_rDA1m (#140556), pAAV-hSyn-DIO-mCherry (#50459-AAV2/9), pAAV-hSyn-hChR2(H134R)-EYFP (#26973-AAV9), pAAV-Syn-ChrimsonR-tdT (#59171-AAV9), pAAV-hSyn-DIO-hM4D(Gi)-mCherry (#44362-AAV2), and pAAV-EF1a-DIO-ChrimsonR-mRuby2-KV2.1-WPRE-SV40 (#124603) were purchased from Addgene. pAAV-FLEX-SaCas9-U6-sgGria1 (#124853), pAAV-FLEX-SaCas9-U6-sgRNA (empty) (#124844) were purchased as plasmids from Addgene and packaged into AAV2/9 at Vector Biolabs. AAV2-Ef1α-DIO-eNpHR3.0-mCherry (Deisseroth) and AAV2-syn-FLEX-ChrimsonR-tdTomato (Boyden) were produced by the Vector Core of the University of North Carolina. AAV9-hsyn-gACh4h(Ach3.8) was purchased from BrainVTA. All viral vectors were stored in aliquots at −80 °C until use.

### Stereotaxic surgery

All stereotaxic surgeries were conducted as described in our animal study protocol using previously described procedures (65) and stereotaxic coordinates (Ma et al., 2020). Mice were first anesthetized with a ketamine (100 mg/kg) plus xylazine (10 mg/kg) solution and placed in a stereotaxic device (AngleTwo, Leica Biosystems). The following stereotaxic coordinates were targeted for viral injections and/or optical fiber implantation: NAc, 1.65 mm from bregma, ±.9 mm lateral from the midline, and -4.6 mm vertical from cortical surface, 0° angle for photometry, 8° angle for optogenetics. PVT, −1.60 mm from bregma, 0.06 mm lateral from midline and −3.30 mm vertical from cortical surface, 6.12° angle for both fiber photometry and optogenetics. VTA, -3.08 mm from bregma, ±.5 mm lateral from the midline, and -4.51 mm vertical from cortical surface. For fiber photometry and optogenetic experiments, an optical fiber (400 μm for photometry, Doric Lenses/Inper; 200 μm for optogenetics, Thorlabs/Inper) was implanted over the target immediately after viral injections and cemented using Metabond Cement System (Parkell) and Jet Brand dental acrylic (Lang Dental Manufacturing). For head fixed training, a head bar was attached with light cure cement (3M RelyX) on top of Metabond. After all surgical procedures, animals were returned to their home cages and placed on a heating pad for 3 hours for post-surgical recovery and monitored for 72 hours with diet supplements. Animals received subcutaneous injections with Metacam (meloxicam, 5 mg/kg) for analgesia and anti-inflammatory purposes. Mice without correct targeting of optical fibers or vectors were excluded from this study.

### Behavior

#### 2-Way Active Avoidance

Mice were trained on the 2AA task replicating previous methodological procedures (21,66) and are detailed below. The behavioral apparatus consisted of a custom-built shuttle box (18 cm × 36 cm × 30 cm) that contained two identical chambers separated by a hurdle (17.5 cm × 6 cm). The hurdle projected 3 cm above the floor and allowed mice easy access to both chambers. The floor consisted of electrifiable metal rods (H10-11M-TC-SF, Coulbourn Instruments) and was connected to a shock generator (H13-15, Coulbourn Instruments). Before each subject was trained/tested, the shuttle box was wiped clean with 70% ethanol. The mouse’s behavior was captured with a USB camera during each session. A speaker located on the top of the shuttle box (50 cm high) was used to deliver the WS. Subjects’ movement and TTLs of WS, US and optogenetic stimulation were recorded and coordinated by ANY-maze (Stoelting).

After a 5-min habituation period, mice were trained with daily sessions of 2AA, each consisting of 30 presentations of the WS (4 kHz, 75 dB, lasting up to 15 s each). Trials in which subjects failed to shuttle to the adjacent chamber before the termination of the WS resulted in the presentation of the unconditioned stimulus (US, 0.6 mA foot shock lasting up to 7 s each) until subjects escaped to the opposite chamber (escape trials). No subject failed to escape the US. For trials in which subjects shuttled to the opposite chamber during the WS, the WS was abruptly terminated, and the US was also prevented (avoidance trials). The inter-trial interval (ITI) was random between 30-40 s for all experiments except Supplementary Fig. 2, where the ITI was increased to 60 s. Avoidance rate was calculated as the percentage of the number of avoidance trials over the total number of trials. For optogenetic and fiber photometry experiments fiber patch cords were attached every session of training. For all photometry experiments, subjects that did not reach 30% avoidance rates by Day 3 were excluded from data analysis. All experiments were conducted in awake, freely moving mice in test chambers during the light cycle.

#### Extinction with response prevention

As shown in Fig. S3, after five days of 2AA training, subjects underwent one session of extinction with response prevention. During this session, animals were confined to a single chamber of the 2AA apparatus using a custom transparent plexiglass divider and presented with 15 WS (15 s each 35-45 s intertrial interval; ITI). On subsequent extinction days, the divider was removed, allowing animals to freely shuttle between chambers while receiving 30 WS presentations (15 s, 30-40 s ITI). No shocks were delivered during extinction, and shuttling did not terminate the WS. To verify that extinction-related changes in photometry signals reflected neural activity rather than photobleaching, a reinstatement session was conducted using the standard 2AA protocol.

#### Head fixed avoidance

The behavioral apparatus comprised a 20 cm diameter foam cylinder connected to a rotary encoder, a head-fixation frame, and a holder for a compressed air nozzle positioned behind the animal, directed toward the base of the tail. The rotary encoder was linked to an interface board connected to the Bpod system (Sanworks, NY). Air puff delivery was controlled through a picospritzer triggered by Bpod TTL output, with pressure set to 50 psi.

Mice were trained daily for 3 days, each session comprising 50 trials. On each trial, a WS (12 kHz, 75 dB, up to 10 s) was presented. If the subject failed to run at least 5 cm on the wheel before WS termination, the US (air puff, up to 5 s) was delivered. If the running requirement was met during the WS, the WS was immediately terminated and the US omitted (avoidance trials). The ITI was 30 s. Avoidance rate was calculated as the percentage of avoidance trials over the total number of trials. Animals typically reached achieved stable performance (≥60% avoidance) by training day 3 or 4.

For the progressive ratio (PR) test, the maximum WS duration was increased to 20 s. The required distance for avoidance started at 25 cm and increased in 5 cm increments. Advancement to the next stage required two consecutive successful avoidance trials. The session ended after six consecutive US deliveries, and the breakpoint was defined as the longest distance at which avoidance was successfully achieved.

### Fiber photometry

Fiber photometry was performed in accordance with previously described methodological procedures (67) and are detailed below. Mice were allowed to habituate to the fiber patch cord in their home cage for approximately 5 min before each behavior test. GCaMP fluorescence and isosbestic autofluorescence signals were excited by the fiber photometry system (Doric Lenses) using two sinusoidally modulated LEDs (473 nm at 211 Hz and 405 nm at 311 Hz) controlled by a standalone driver (DC4100, ThorLabs). Both LEDs were combined via a commercial Mini Cube fiber photometry apparatus (Doric Lenses) into a fiber patch cord (400-µm core, 0.48 NA) connected to the brain implant in each mouse. The light intensity at the interface between the fiber tip and the animal was adjusted from 10 µW to 20 µW (but was constant throughout each test session for each mouse). An RZ5P fiber photometry acquisition system with Synapse software (Tucker-Davis Technologies) collected and saved real-time demodulated emission signals and behavior-relevant TTL inputs. For each trial, signals (F473 nm) were compared with autofluorescence signals (F405 nm) to control for movement and bleaching artifacts. Signal data were de-trended by first applying a least-squares linear fit to produce F_fitted 405 nm_, and dF/F was calculated as (F_473 nm_ – F_fitted 405 nm_)/F_fitted 405 nm_. All signal data are presented as the z-score of the dF/F from baseline (pre-WS/pre-Event) segments. In Supplementary Fig. 6, a third channel was added (560 nm LED driven at 711 Hz). This signal was compared to autofluorescence signals (F405 nm) as described.

For Fig. 3 and S5, an RZ10x fiber photometry acquisition system (Tucker-Davis Technologies) with integrated drivers, LEDs (405nm, 465nm, 560nm) and photosensors was used to collect and demodulate emission signals with analogous protocols and analysis.

#### Fiber photometry with optogenetic and chemogenetic manipulations

In Fig. 2, mice with dLight in NAc unilaterally and halorhodopsin or mCherry in pPVT underwent 2AA training. Fiber photometry recordings of dLight were taken during training on days 1, 3, and 5. In addition, a red light (Opto Engine 635nm) was triggered at the completion of all avoidance trials concurrent with WS offset for a 5 s duration.

In Fig. 3K-O, mice were injected with dLight in NAc, DREADDs (Gi) in NAc ChAT+ neurons, and ChrimsonR or mCherry in pPVT. Fiber photometry recordings of dLight were taken in a novel context on separate days during optogenetic stimulation (60s, 20Hz, 10 trials) of pPVT terminals. For within mouse comparisons, each subject was given a saline injection I.P. 30 minutes prior to recording. The following day the same subjects were given a CNO injection (5 mg/kg) I.P. 30 minutes prior to recording. The mice were training in 2AA for 5 days before repeating saline and CNO recording conditions.

### Optogenetics

In Fig. 1, 2 and S2, mice were subjected to five 2AA sessions (one session per day), on training days 1-5 a yellow light (Ce:YAG + LED Driver, Doric) was triggered at the completion of all avoidance trials concurrent with WS offset for a 5 s duration. For experiments with halorhodopsin stimulation, the light was powered on for the 5 seconds and light intensity at the interface between the fiber tip and the implant cannula was ∼10 mW. For ChrimsonR experiments, the stimulation was ∼10 mW, 20 Hz, 20% duty cycle.

### Electrophysiology

Mice were trained in 2AA for 5 days, and two hours following the last session, the subjects were then sacrificed for slice recordings by an experimenter blind to conditions. For yoked controls, each was paired with a trained subject. The pairs were trained concurrently wherein the yoked subject received the same duration of both WS and shock.

Acute brain slices were prepared according to previously described protocols (65,66,68). Mice were anesthetized with isoflurane and transcardially perfused with an ice-cold NMDG cutting solution containing (in mM): 92 mM N-Methyl-D-glucamine, 2.5 mM KCl, 1.25 mM NaH2PO4, 10 mM MgSO4, 0.5 mM CaCl2, 30 mM NaHCO3, 20 mM glucose, 20 mM HEPES, 2 mM thiourea, 5 mM Na-ascorbate, 3 mM Na-pyruvate, at 7.3-7.4 pH, saturated by 95% O2 and 5% CO2. Coronal sections (300 µm thickness) containing the NAc were cut in the ice-cold NMDG cutting solution using a VT1200S automated vibrating-blade microtome (Leica Biosystems), and were subsequently transferred to a heated incubation chamber containing the NMDG cutting solution at 34-35 °C. After recovery for 12 min, slices were transferred to a room temperature (20-24 °C) holding chamber containing a HEPES-modified artificial cerebrospinal fluid (92 mM NaCl, 2.5 mM KCl, 1.25 mM NaH2PO4, 2 mM MgSO4, 2 mM CaCl2, 30 mM NaHCO3, 25 mM glucose, 20 mM HEPES, 2 mM thiourea, 5 mM Na-ascorbate, 3 mM Na-pyruvate, at 7.3 pH, saturated by 95% O2 and 5% CO2) and remained in the holding chamber until recordings.

Whole-cell patch-clamp recordings were conducted under visual guidance using an Olympus BX51 microscope with transmitted light illumination, and ChAT+ neurons were identified based on their fluorescence (mCherry). Recordings from ChAT+ neurons in the NAc were obtained with Multiclamp 700B amplifiers (Molecular Devices) and data were filtered at 1-2 kHz using a 1440A Digidata Digitizer and Clampex software (Molecular Devices). For recordings, slices were transferred to the recording chamber and constantly supplied with a room-temperature ACSF (118 mM NaCl, 2.5 mM KCl, 26.2 mM NaHCO3, 1 mM NaH2PO4, 20 mM glucose, 2 mM MgCl2, and 2 mM CaCl2, at pH 7.4, saturated by 95% O2 and 5% CO2, flow rate = 2.0-2.5ml/min). Pharmacological agents were added to the ACSF and bath applied. All recordings were made with borosilicate glass pipettes with tip resistance of 3-6 MΩ. pipettes were filled with Cs-based internal solution containing (in mM): 117 mM Cs methanesulfonate, 10 mM HEPES, 2.5 mM MgCl2, 2 mM Na2-ATP, 0.4 mM Na2-GTP, 10 mM Na2-phosphocreatine, 0.6 mM EGTA, 5 mM QX-314 at pH 7.2 and 288-290 mOSM. Optogenetically evoked synaptic responses were achieved by illuminating the acute slices with blue LED (470 nm, CoolLED) drive ChR2-expressing terminals. Light stimulation was delivered every 15 seconds and synaptic responses were recorded at holding potentials of −60 mV for AMPA-receptor-mediated responses and +40 mV for NMDA-receptor-mediated responses. NMDA-receptor-mediated responses were quantified as the EPSC magnitude at +40 mV (50 ms following stimulation). Evoked EPSCs were recorded with picrotoxin (100 μM) added to the ACSF. To assess presynaptic function, a paired-pulse stimulation protocol (50 ms inter-stimulus interval) was used to evoke double-EPSCs, and the paired-pulse ratio (PPR) was quantified as the ratio of the peak amplitude of the second EPSC to that of the first EPSC. Rise time and decay time of EPSCs were measured between 20–80% of amplitude, respectively. For the measurement of rectification of AMPA current, spermine (100 μM) was added in the internal solution, while picrotoxin (100 μM) and NMDA receptor blocker D-(-)-2-Amino-5-phosphonopentanoic acid (AP-5) (50 μM) were added in ACSF. Neurons were held at −70, −40, 0, +40 or +70 mV, respectively. The rectification index (RI) was calculated by dividing the absolute amplitude of average EPSC at +40 mV by that at −70 mV. Naspm trihydrochloride (NASPM, 100 µM, Tocris) was applied to the slice while stimulating every 15 s. A stable baseline was recorded for at least 5 min followed by a 10 min application of NASPM. The average amplitude of the EPSC was calculated during the 5 min period prior to NASPM application (baseline) and 10 min after drug application, with the EPSC amplitude normalized to the baseline period.

### Fast-Scan Cyclic Voltammetry

Fast-scan cyclic voltammetry was performed as previously described (32). Carbon-fiber electrodes (CFEs) were prepared with a cylindrical carbon-fiber (7mm diameter, ∼150mm of exposed fiber) inserted into a glass pipette, and were conditioned with an 8-ms-long triangular voltage ramp (-0.4 to 1.2 and back to -.4 V vs Ag/AgCl reference at 400 V/s) delivered every 15 ms. CFEs showing current >1.8 mA or <1.0 mA in response to the voltage ramp at ∼0.6 V were discarded. During recording, CFEs were held at -0.4 V versus Ag/AgCl, and a triangular voltage ramp was delivered every 100 ms. DA signals were evoked by single pulse optical stimulations, where a fiberoptic (200mm diameter, 0.22 NA, ThorLabs) connected to a blue LED (470 nm, 1.8 mW, ThorLabs) was placed over the carbon fiber and light pulses (∼0.6 ms) were delivered every 2 min. Data were collected with a retrofit headstage (CB-7B/ EC with 5 MX resistor) using a Multiclamp 700B amplifier (Molecular Devices) after low-pass filter at 3 kHz and digitized at 100 kHz using a DA board (NI USB-6229 BNC, National Instruments). Data acquisition and analysis were performed using a custom-written software, VIGOR, in Igor Pro (Wavemetrics) using mafPC (courtesy of MA Xu-Friedman). The current peak amplitudes of the evoked DA transients were converted to DA concentration according to the post experimental calibration using 1-3 mM DA.

### Histology and immunofluorescence

Animals were deeply anesthetized with euthanasia solution (Vet One) and transcardially perfused with PBS (pH 7.4, 4 °C), followed by paraformaldehyde (PFA) solution (4% in PBS, 4 °C). After extraction, brains were post-fixed in 4% PFA at 4 °C for a minimum of 2 h and subsequently cryoprotected by transferring to a 30% PBS-buffered sucrose solution until brains were saturated (for over 24 h). Coronal brain sections (50 μm) were cut using a freezing microtome (SM 2010R, Leica).

#### Sample preparation and ISH procedure for RNAscope

For Fig. 5 and S7, Fresh-frozen brains from adult male C57BL/6NJ mice (8–12 weeks old) were sectioned at a thickness of 16 µm using a Cryostat (Leica Biosystems). Sections were collected onto Superfrost Plus glass slides (Daigger Scientific), immediately placed on dry ice and subsequently transferred to a −80 °C freezer. mRNA signal for *Chat* and *saCas9* was detected using the RNAscope fluorescent kit (Advanced Cell Diagnostics). Specifically, glass slides with sections spanning the entire anteroposterior spread of the PVT were removed from the −80 °C freezer, fixed with freshly prepared ice-chilled 4% PFA for 15 min at 4 °C and then dehydrated using a series of ethanol solutions at different concentrations (5 min each, room temperature): 1 X 50%, 1 X 70% and 2 X 100%. Next, sections were treated with Protease IV (Advanced Cell Diagnostics) at room temperature for 30 min. Slides were then washed with PBS twice (1 min each) and dried for 5 min at room temperature, and sections were circled with an ImmEdge Hydrophobic Barrier PAP Pen (Vector Laboratories). Hybridization was performed on a HybEZ oven for 2 h at 40 °C using a *Chat* and *saCas9* probe (Advanced Cell Diagnostics). After this, the slides were washed twice with washing buffer (2 min each), then incubated with Hybridize Amp 1-FL for 30 min, Hybridize Amp 2-FL for 15 min, Hybridize Amp 3-FL for 30 min and Hybridize Amp 4-FL for 15 min. Next, the slides were washed twice with washing buffer (2 min each) and coverslips added using Diamond Prolong antifade mounting medium with DAPI (Thermo Fisher Scientific).

### Data analysis and statistics

#### Statistical analyses

All data were plotted and analyzed with GraphPad Prism, with Python, Matlab, or Clampfit (Molecular Devices). A summary table of statistical results is provided (Supplemental Table 1).

Standard statistical tests used are described in figure legends and statistics table. All data was tested for normality with Shapiro-Wilk tests and non-parametric or mixed models were used if non-passing. Unbalanced area under the curve datasets were analyzed using linear mixed-effects models (REML) with a random intercept for subject to account for repeated measures using statsmodels mixedlm in Python. Main effects of categorical factors were tested using F tests on model-estimated contrasts. Pairwise comparisons were performed using t tests on contrasts and p-values adjusted with the Sidak method for multiple comparisons. Balanced datasets were tested with mixed-effects models (REML) in GraphPad Prism.

#### RL Model

A reinforcement learning model (69) was fit to the sample mean using Matlab:

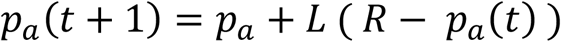

where pa is the probability of avoidance, t is the trial, L is the learning rate, and R is the set value of avoidance. L, R, and a starting pa were fit as free parameters using least sum of squares. Standard error was estimated within groups using a 1000 bootstrap permutation test with replacement. A 5000 bootstrap permutation test with a null distribution generated by randomly reassigning group labels was used to assess the significance of differences between group parameters. The simulation in Fig. 1K was created using extreme high and low R and L parameters seen in real datasets. Trial outcome was generated using softmax probability (β=.8) and 100 simulations were averaged.

#### Bilinear Model

Where R and L were found to be highly correlated (Fig. S2), a model where “rate of learning” and “asymptote of behavior” were more directly estimated was used. A bilinear model was fit to sample mean avoidance rate, that consisted of one sloping line fit to the data intersecting with one horizontal line. The best fit was found as the intersection point with the lowest sum of squares error to the data and the statistical difference between groups of the corresponding slope, intersection point, and horizontal y values was tested with 1000 bootstrap permutation test with a null distribution generated by randomly reassigning group labels. Standard error was estimated within groups using a 1000 bootstrap permutation test with replacement.

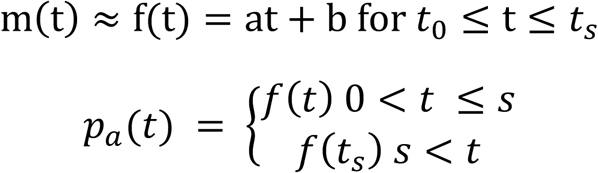

t being the trial number, m the mean of the sample data, s being the intersection point trial, and f(t) a linear fit using polyfit from the numpy package.

### Schematics

Created in BioRender. https://BioRender.com/ke4lom1

## Acknowledgements

The authors thank the members of the Section on the Neural Circuits of Emotion and Motivation and the Laboratory on Neural Circuits and Behavior of NIMH who offered scientific feedback. In addition, we thank the NIMH IRP Rodent Behavioral Core and the NIMH IRP Systems Neuroscience Imaging Resource for supporting this work. Finally, we thank Dr. Yulong Li for providing the GRAB ACh3.8 sensor and Dr. Michael Halassa for offering scientific and writing feedback.

## Author contributions

Conceptualization: E.E.M. and M.A.P.; Data acquisition; E.E.M., J.M., D.L., K.Y., M.E.A., Y.L., H.C.G.; Data analysis: E.E.M., D.L., K.Y., M.E.A.; Project management and supervision; M.A.P., B.B.A., V.A.A.; Writing original draft: E.E.M, M.A.P.

