## Supplementary Figures and Table for "A synaptic mechanism for encoding the learned value of action-derived safety"

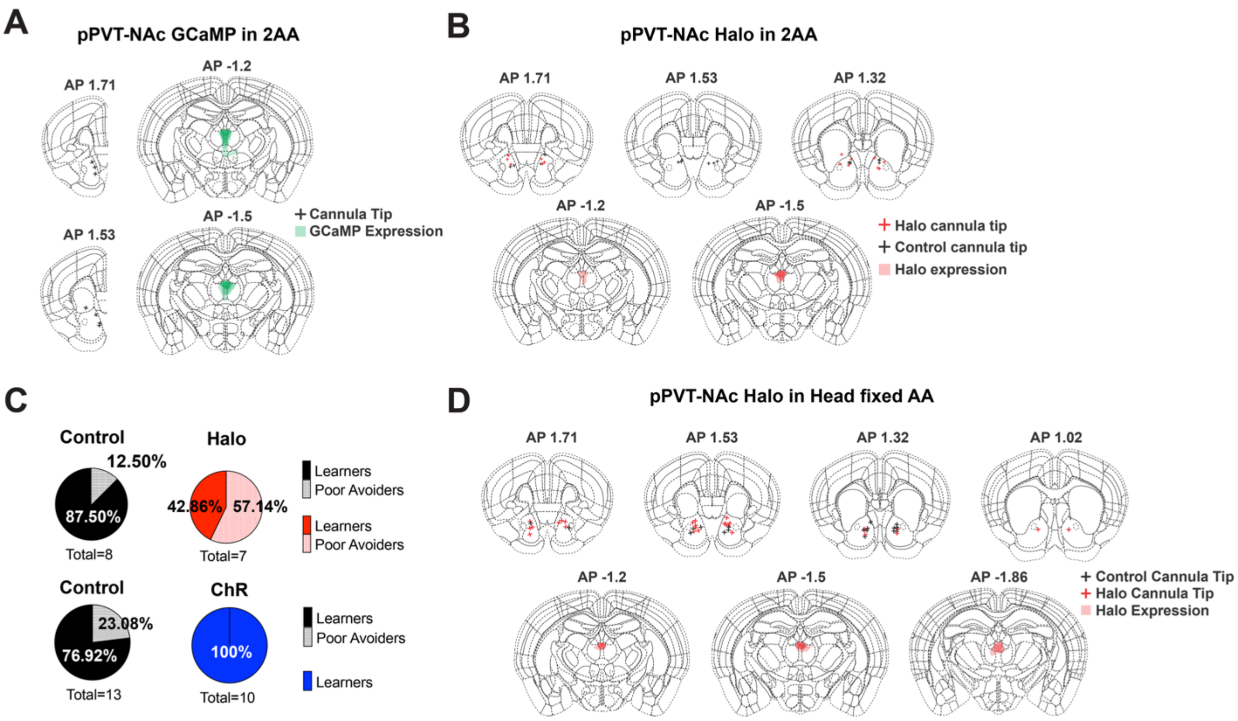


**Supplementary Figure 1: Related to Figure 1. (A)** Map of optic fiber cannula tip locations for photometry experiment in figures 1A-G (left). Map of viral spread of GCaMP at bregma -1.2 and -1.5 (right). **(B)** Map of optic fiber cannula tip locations for optogenetics experiment in figures 1H-L (top). Map of viral spread of halorhodopsin (Halo) at bregma -1.2 and -1.5 (bottom). **(C)** Proportion of poor avoiders (<30% avoidance day 3 of training) per optogenetic experimental group in figures 1H-L and S3. **(D)** Map of optic fiber cannula tip locations for optogenetics experiment in figures 1M-O (top). Map of viral spread of halorhodopsin at bregma -1.2 and -1.5 (bottom).

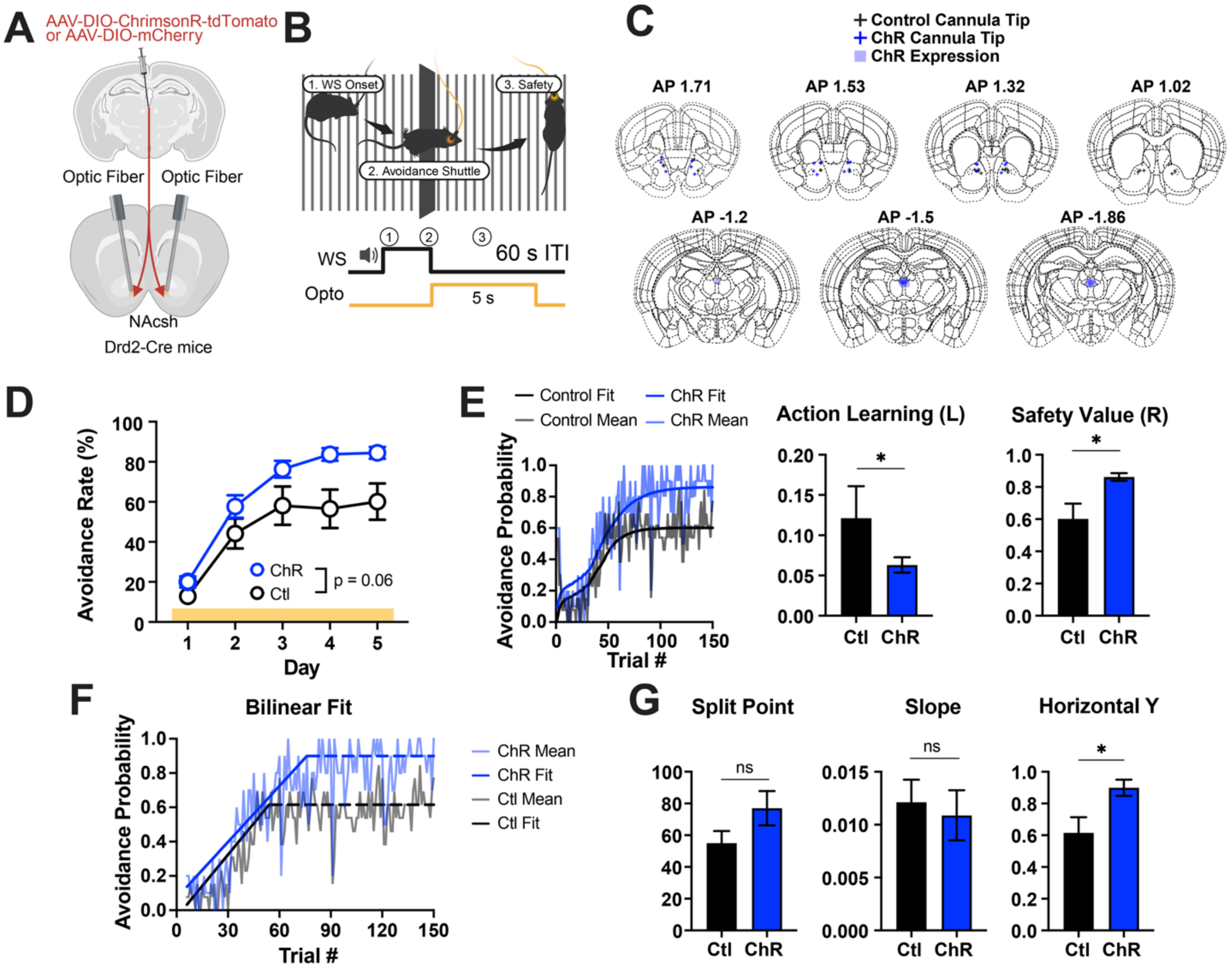


**Supplementary Figure 2: pPVT-NAc activation boosts safety value. (A)** Drd2-Cre mice were injected with ChrimsonR (ChR) or fluorescent control in pPVT to bilateral stimulate terminals in NAc. **(B)** Schematic of closed-loop optogenetic manipulation in the 2AA task (see Methods). **(C)** Map of optic fiber cannula tip locations (top). Map of viral spread of ChrimsonR at bregma -1.2, -1.5, and -1.86 (bottom). **(D)** Performance in the 2AA task with excitation of pPVT-NAc at safety (Control n=13, ChR n=10 mice). 2-way Anova group effect: p=0.0554. **(E)** Avoidance probability across trial with best fit Rescorla-Wagner model probability by group (left). Best fit learning and value parameters by group with bootstrap estimated error (middle, right). Bootstrap permutation test for group difference: Learning p=0.0412; Value p=0.0208. **(F)** Avoidance probability across trial with best fit bilinear model by group. **(G)** Bilinear model best fit parameters by group with bootstrap estimated error. Bootstrap permutation test for group difference: Split point p=0.084; Slope p=0.7284; Horizontal p=0.0188. Data are shown as mean ± s.e.m. *p < 0.05, ns: p > 0.05.

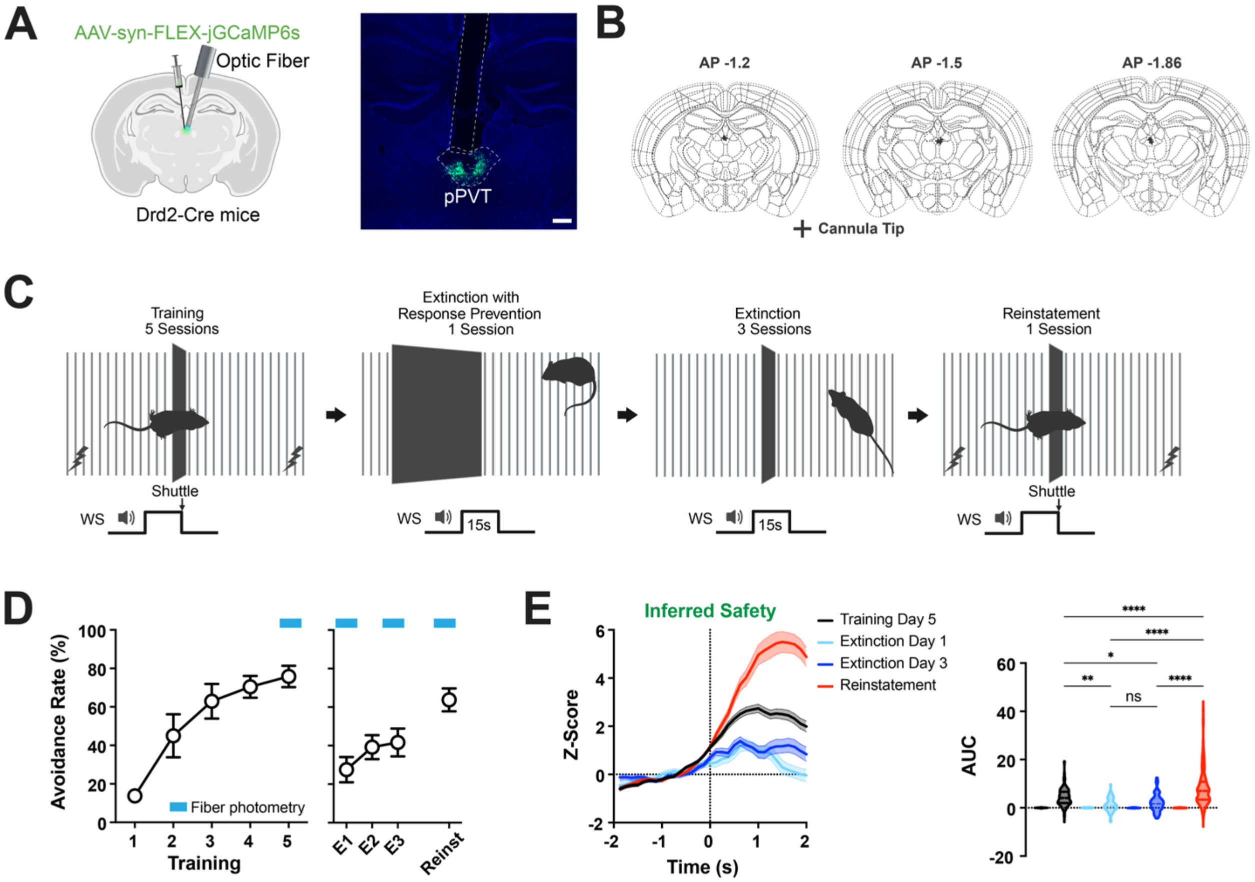


**Supplementary Figure 3: pPVT tracks safety devaluation. (A)** Drd2-Cre mice were injected with GCaMP to record during 2AA and extinction (left). Representative image of injection and recording site (right). Scale bar, 200um. **(B)** Map of optic fiber cannula tip locations. **(C)** Schematic of training protocol (see Methods). **(D)** Performance in the 2AA task (n=8 mice). **(E)** GCaMP dF/F z-score normalized at shuttle (safety) with optogenetic inhibition by day of training (left). Quantification of post event activity (right). AUC, pairwise comparisons between groups, linear mixed-effects model for repeated measures: see statistics table. Data are shown as mean ± s.e.m. $^*p < 0.05$, $^{**}p < 0.01$, $^{****}p < 0.0001$, ns: $p > 0.05$.

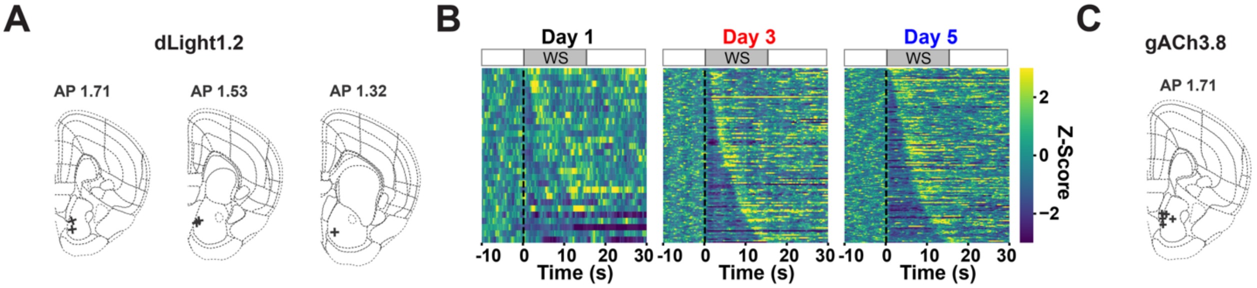


**Supplementary Figure 4: Related to figures 2 and 3. (A)** Map of fiber optic cannula tip location for dLight photometry experiment in figures 2A-F. **(B)** Heatmaps of dLight signals during avoidance trials, ordered by avoidance latency (low to high). Related to figures 2A-F. **(C)** Map of fiber optic cannula tip location for gACh photometry experiment in figures 3D-J.

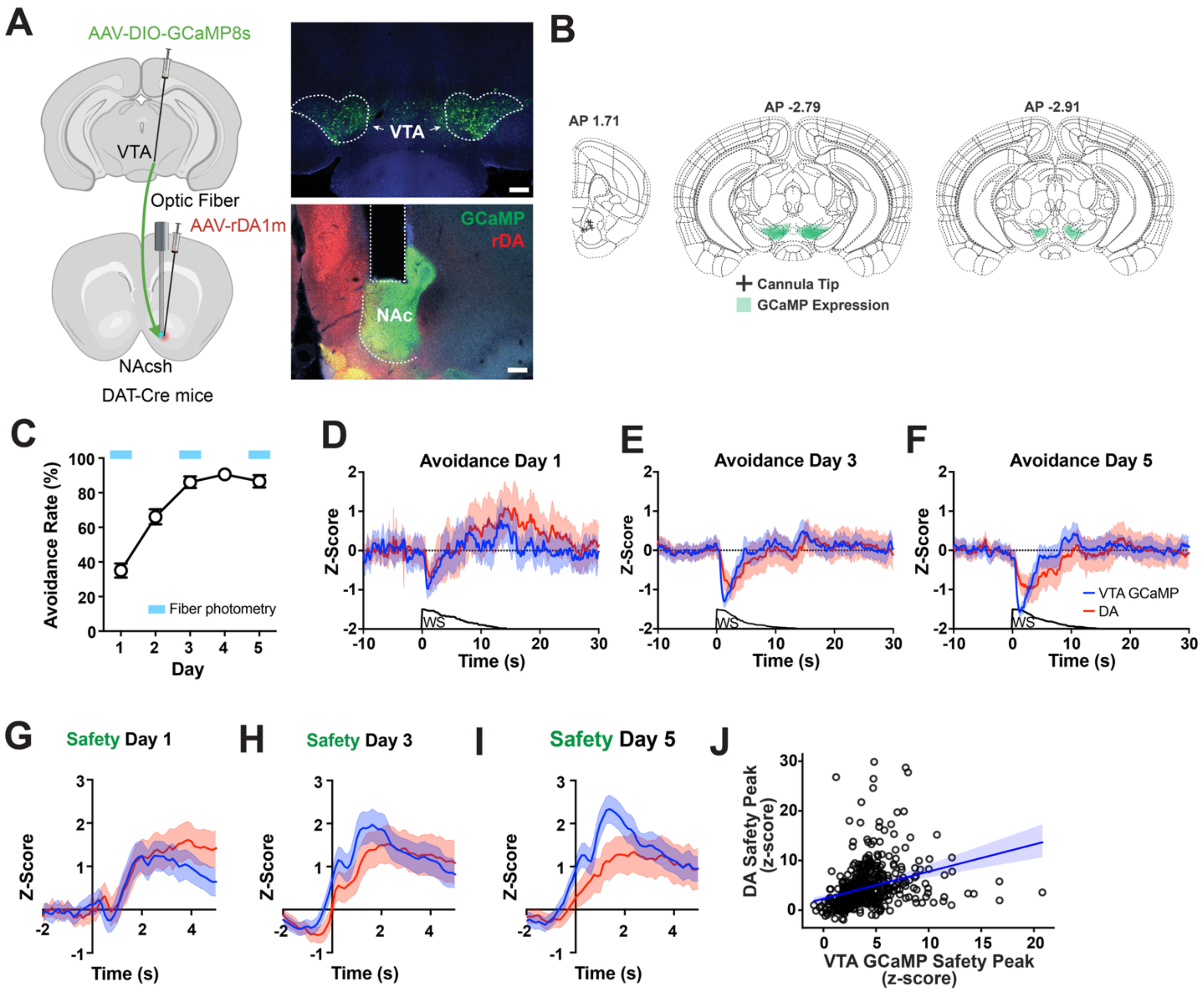


**Supplementary Figure 5: rDA1m sensor activity matches dopaminergic terminals. (A)** DAT-Cre mice were injected with GCaMP8s in the VTA and rDA1m in the NAc to validate the GRAB DA sensor (left). Representative images of injection and recording sites (right). Scale bars, 200um. **(B)** Map of optic fiber cannula locations (left) and GCaMP viral expression in VTA at bregma -2.79 and -2.91 (right). **(C)** Performance in the 2AA task (n=6 mice). **(D-F)** GCaMP (blue) and rDA (red) dF/F z-score normalized in all avoidance trials on training days 1 (**D**), 3 (**E**) and 5 (**F**), with average WS duration below. **(G-I)** dF/F z-score normalized at safety on training days 1 (**G**), 3 (**H**) and 5 (**I**). **(J)** Correlation between peak z-score value at safety by trial of GCaMP and rDA. Pearson correlation: $p<0.0001$. Data are shown as mean ± s.e.m.

## A

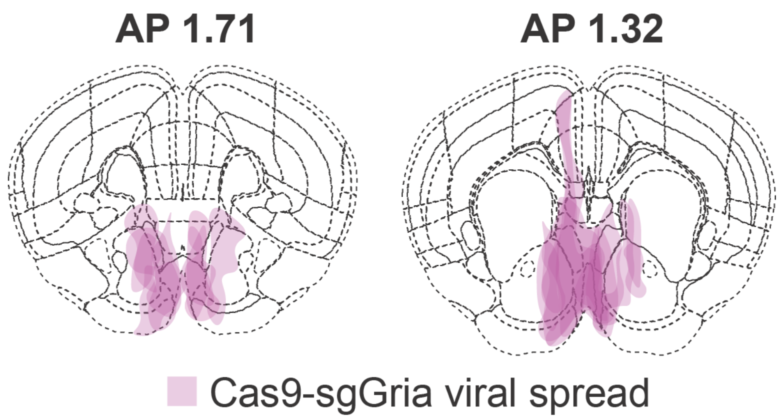


**Supplementary Figure 6: Related to figure 5. (A)** Map of viral spread for sgGria KO in figures 5N-Q at bregma 1.71 and 1.32.

**Supplementary Table 1: Statistics Table**

| Figure | Comparison | Statistical Test | P Value |
|---|---|---|---|
| **Figure 1F** | Shuttle Initiation AUC Day Effect | AUC~C(Day)+(1\|Subject) F(2, 285) = 3.512 | * p=0.0311 |
| | Day 1 vs Day 3 | Pairwise comparison Sidak corrected | p=0.9937 |
| | Day 1 vs Day 5 | Pairwise comparison Sidak corrected | p=0.3913 |
| | Day 3 vs Day 5 | Pairwise comparison Sidak corrected | * p=0.0324 |
| **Figure 1G** | Safety AUC Day Effect | AUC~C(Day)+(1\|Subject) F(2, 398) = 5.700 | * p=0.0036 |
| | Day 1 vs Day 3 | Pairwise comparison Sidak corrected | p=0.7459 |
| | Day 1 vs Day 5 | Pairwise comparison Sidak corrected | * p=0.0209 |
| | Day 3 vs Day 5 | Pairwise comparison Sidak corrected | * p=0.0161 |
| **Figure 1J** | Day x Group | 2 Way ANOVA F (5, 65) = 1.780 | p=0.1294 |
| | Day | F (2.758, 35.86) = 13.16 | **** p<0.0001 |
| | Group | F (1, 13) = 5.573 | * p=0.0345 |
| **Figure 1L** | Learning Ctl vs Halo | Bootstrap permutation test LCtl-LHalo=-0.096341, 95% CI5000=[-0.12019125,0.1031405] | p=0.1032 |
| | Value Ctl vs Halo | Bootstrap permutation test RCtl-RHalo=0.34851, 95% CI5000=[-0.33509075,0.3382775] | * p=0.0448 |
| **Figure 1N** | First day of training avoidance | Unpaired t test t=1.387, df=26 | p=0.1771 |
| | Last day of training avoidance | Unpaired t test t=1.178, df=26 | p=0.2493 |
| **Figure 1O** | Ctl Laser Off vs On | Paired t test t=0.7042, df=12 | p=0.4947 |
| | Halo Laser Off vs On | Paired t test t=3.458, df=15 | ** p=0.0035 |
| **Figure 2E** | Shuttle Initiation AUC Day Effect | AUC~C(Day)+(1\|Subject) F(2, 302) = 2.001 | p=0.1370 |
| **Figure 2F** | Safety AUC Day Effect | AUC~C(Day)+(1\|Subject) F(2, 201) = 1.929 | p=0.1479 |
| **Figure 2I** | Day x Group | 2 Way ANOVA F (8, 52) = 0.3850 | p=0.9238 |
| | Day | F (2.094, 27.22) = 57.84 | **** p<0.0001 |
| | Group | F (2, 13) = 0.6944 | p=0.517 |
| **Figure 2K** | Day | AUC~Day*Group+(1\|Subj) F(2, 574) = 10.488 | **** p<0.0001 |
| | Group | F(1, 574) = 4.742 | * p=0.0298 |
| | Day x Group | F(2, 574) = 0.457 | p=0.6336 |

| | | | |
|---|---|---|---|
| | Ctl vs Halo Day 1 | Pairwise comparison Sidak corrected | p=0.0857 |
| | Ctl vs Halo Day 3 | Pairwise comparison Sidak corrected | ** p=0.0075 |
| | Ctl vs Halo Day 5 | Pairwise comparison Sidak corrected | * p=0.0259 |
| **Figure 3C** | DA Amplitude Baseline vs DHBE | Paired t test t=24.68, df=4 | **** p<0.0001 |
| **FIgure3I** | Shuttle Initiation AUC Day Effect | AUC~C(Day)+(1\|Subject) F(2, 217) = 4.073 | * p=0.0183 |
| | Day 1 vs Day 3 | Pairwise comparison Sidak corrected | p=0.4551 |
| | Day 1 vs Day 5 | Pairwise comparison Sidak corrected | * p=0.0182 |
| | Day 3 vs Day 5 | Pairwise comparison Sidak corrected | p=0.2072 |
| **Figure 3J** | Safety AUC Day Effect | AUC~C(Day)+(1\|Subject) F(2, 345) = 5.337 | ** p=0.0052 |
| | Day 1 vs Day 3 | Pairwise comparison Sidak corrected | p=0.1370 |
| | Day 1 vs Day 5 | Pairwise comparison Sidak corrected | ** p=0.0036 |
| | Day 3 vs Day 5 | Pairwise comparison Sidak corrected | p=0.2902 |
| **Figure 3O** | Training | AUC~Day*Treatment*Time + (1\|Subject) F(1, 312) = 5.129 | * p=0.0242 |
| | Treatment | F(1, 312) = 14.011 | *** p=0.0002 |
| | Time | F(1, 312) = 1.126 | p=0.2894 |
| | Treatment x Time | F(1, 312) = 7.007 | ** p=0.0085 |
| | Training x Time | F(1, 312) = 2.557 | p=0.1108 |
| | Treatment x Training | F(1, 312) = 3.084 | p=0.0800 |
| | Treatment x Training x Time | F(1, 312) = 1.545 | p=0.2149 |
| | Pre vs On Pre-Training Saline | Pairwise comparison Sidak corrected | p=0.3277 |
| | Pre vs On Pre-Training CNO | Pairwise comparison Sidak corrected | ** p=0.0089 |
| | Pre vs On Post-Training Saline | Pairwise comparison Sidak corrected | p=0.0708 |
| | Pre vs On Post-training CNO | Pairwise comparison Sidak corrected | p=0.9668 |
| | On Pre-Training Saline vs Pre-Training CNO | Pairwise comparison Sidak corrected | p=0.9027 |
| | On Pre-Training Saline vs Post-Training Saline | Pairwise comparison Sidak corrected | **** p<0.0001 |
| | On Pre-Training Saline vs Post-Training CNO | Pairwise comparison Sidak corrected | p=0.9772 |
| | On Pre-Training CNO vs Post-Training Saline | Pairwise comparison Sidak corrected | **** p<0.0001 |

| | | | |
|---|---|---|---|
| | On Pre-Training CNO vs Post-Training CNO | Pairwise comparison Sidak corrected | p=0.2119 |
| | On Post-Training Saline vs Post-Training CNO | Pairwise comparison Sidak corrected | ** p=0.0018 |
| **Figure 4E** | AMPA Currents | Unpaired t test t=3.241, df=39 | ** p=0.0024 |
| **Figure 4F** | AMPA Rise Time | Unpaired t test t=2.683, df=39 | * p=0.0107 |
| **Figure 4G** | AMPA Decay Time | Unpaired t test t=3.193, df=39 | ** p=0.0028 |
| **Figure 4H** | NMDA Currents | Unpaired t test t=1.446, df=39 | p=0.1562 |
| **Figure 4I** | AMPA/NMDA Ratio | KS normality test: yoked, trained | ** p=0.002, *** p=0.0006 |
| | AMPA/NMDA Ratio | Mann Whitney test U=118 | * p=0.0168 |
| **Figure 4J** | Paired-pulse Ratio | Unpaired t test t=3.775, df=39 | *** p=0.0005 |
| **Figure 4K** | Input x Treatment | 2way ANOVA F (4, 76) = 7.266 | *** p<0.0001 |
| | Input | F (4, 76) = 701.3 | *** p<0.0001 |
| | Treatment | F (1, 19) = 8.625 | ** p=0.0085 |
| | 70mV | Bonferroni's multiple comparisons test | **** p<0.0001 |
| | Rectification Index | Unpaired t test t=2.257, df=19 | * p=0.0360 |
| **Figure 4L** | Inhibition by NASPM | Unpaired t test t=5.545, df=10 | *** p=0.0002 |
| **Figure 5E** | AMPA Currents | Unpaired t test t=4.257, df=22 | *** p=0.0003 |
| **Figure 5F** | AMPA Rise Time | Unpaired t test t=2.494, df=22 | * p=0.0206 |
| **Figure 5G** | AMPA Decay Time | Unpaired t test t=2.288, df=22 | * p=0.0321 |
| **Figure 5H** | Paired-pulse Ratio | Unpaired t test t=3.825, df=22 | *** p=0.0009 |
| **Figure 5J** | NMDA Currents | Unpaired t test t=0.7933, df=19 | p=0.4374 |
| **Figure 5K** | AMPA/NMDA Ratio | Unpaired t test t=2.421, df=19 | * p=0.0256 |
| **Figure 5M** | Inhibition by NASPM | Unpaired t test t=3.173, df=9 | * p=0.0113 |
| **Figure 5O** | Day x Group | Mixed effects F (4, 40) = 2.140 | p=0.0936 |
| | Day | F (1.645, 16.45) = 52.67 | **** p<0.0001 |
| | Group | F (1, 10) = 5.082 | * p=0.0478 |
| **Figure 5Q** | Learning Empty vs sgGria1 | Bootstrap permutation test LCtl-LHalo=-0.0137, 95% CI5000=[-0.01905325,0.01904415] | p=0.2136 |
| | Value Empty vs sgGria1 | Bootstrap permutation test RCtl-RHalo=0.1512, 95% CI5000=[-0.1499,0.1458] | * p=0.0372 |
| **Figure S2D** | Day x Group | 2way ANOVA F (5, 105) = 1.987 | p=0.0865 |
| | Day | F (2.631, 55.24) = 77.02 | **** p<0.0001 |
| | Group | F (1, 21) = 4.116 | p=0.0554 |
| **Figure S2E** | Learning Ctl vs ChR | Bootstrap permutation test LCtl-LChR=0.05827, 95% CI5000=[-0.0645,0.0565] | * p=0.0412 |

| | | | |
|---|---|---|---|
| | Value Ctl vs ChR | Bootstrap permutation test RCtl-RChR=-0.26011, 95% CI5000=[-0.2347,0.2493] | * p=0.0208 |
| **Figure S2G** | Split point | Bootstrap permutation test sCtl-sChR=-22.0, 95% CI5000=[-25.0,21.0] | p=0.084 |
| | Slope | Bootstrap permutation test aCtl-aChR=0.001246, 95% CI5000=[-0.0060,0.0064] | p=0.7284 |
| | Horizontal Y | Bootstrap permutation test f(ts)Ctl-f(ts)ChR=-0.283716, 95% CI5000=[-0.1943,0.2385] | * p=0.0188 |
| **Figure S3E** | Shuttle AUC Day Effect | AUC~C(Day)+(1\|Subject) F(3.0, 490) = 29.413 | **** p<0.0001 |
| | TD5 vs. ExD1 | Pairwise comparison Sidak corrected | ** p=0.0031 |
| | TD5 vs. ExD3 | Pairwise comparison Sidak corrected | * p=0.0237 |
| | TD5 vs. Reinst | Pairwise comparison Sidak corrected | **** p<0.0001 |
| | ExD1 vs. ExD3 | Pairwise comparison Sidak corrected | p=0.9421 |
| | ExD1 vs. Reinst | Pairwise comparison Sidak corrected | **** p<0.0001 |
| | ExD3 vs. Reinst | Pairwise comparison Sidak corrected | **** p<0.0001 |
| **Figure S5I** | Safety Peak GCaMP vs DA | Pearson r=0.3160 | **** p<0.0001 |
